# A framework for linking hemispheric, full annual cycle prioritizations to local conservation actions for migratory birds

**DOI:** 10.1101/2023.04.03.534556

**Authors:** William V. DeLuca, Nathaniel E. Seavy, Joanna Grand, Jorge Velásquez-Tibatá, Lotem Taylor, Cat Bowler, Jill L. Deppe, Erika J. Knight, Gloria M. Lentijo, Timothy D. Meehan, Nicole L. Michel, Sarah P. Saunders, Nolan Schillerstrom, Melanie A. Smith, Chad Witko, Chad B. Wilsey

## Abstract

The conservation of migratory birds poses a fundamental challenge: their conservation requires coordinated action across the hemisphere, but those actions must be designed and implemented locally. To address this challenge, we describe a multi-level framework for linking broad-scale, full annual cycle prioritizations to local conservation actions for migratory birds. We developed hemisphere-scale spatial prioritizations for the full annual cycle of migratory birds that breed in six different ecosystems in North America. The full annual cycle prioritizations provide a hemispheric context within which regional priorities can be identifieed. Finer resolution, regional prioritizations can then inform local conservation actions more effectively. We describe the importance of local conservation practitioner contributions at each level of the process and provide two examples of regional spatial prioritizations that were developed to guide local action. The fierst example focused on coastal North and South Carolina, USA, and used information on marsh birds, shorebirds, ecological integrity, and co-benefiets for people to identify Cape Romain, South Carolina as a high-priority site for conservation action. The second example in Colombia used information on migrant and resident birds to identify the Cauca Valley as a high priority site. The multi-level conceptual framework we describe is one pathway for identifying sites for implementation of local conservation actions that are guided by conservation priorities for migratory birds across their full annual cycle.

## INTRODUCTION

Reversing the alarming decline of migratory birds breeding in North America (Rosenberg et al., 2019; Wilcove & Wikelski, 2008) requires approaches that span hemispheres, making their conservation notoriously challenging (Runge, Martin, Possingham, Willis, & Fuller, 2014). Strategies for the conservation of migratory birds must include information across the full annual cycle of a species, often spanning an entire hemisphere, while guiding the implementation of conservation actions that are relevant and appropriate within a local context. For decades, ecologists have used the concept of multi-scale habitat selection as a framework to explain and study the ability of animals to successfully move across broad spatial extents (e.g., McGarigal et al. 2016). Similarly, the recognition that a multi-level perspective is valuable has also been used to explain the challenges facing conservation and environmental management (Bodin, Crona, Thyresson, Golz, & Tengö, 2014; Cash et al., 2006; Erasmus, Freitag, Gaston, Erasmus, & Jaarsveld, 1999).

Although multi-level strategies have been used in some conservation planning efforts (e.g., Gonthier et al. 2014; Lagabrielle et al. 2018), most of these approaches have focused on levels of ecological organization. Perhaps the best example of this is the coarse fielter/fiene fielter approach developed by The Nature Conservancy to ensure that conservation planning considers both ecological communities (coarse fielter) and individual species (fiene fielter; Noss, 1987). However, examples of applying multi-level approaches to migratory bird conservation have been limited (however, see Palumbo et al. 2021). The limited application of these approaches to migratory bird conservation may in part be due to a lack of necessary ecological data across the annual cycle as well as the complexity of bringing together diverse partners who can represent multiple scales of decision making (Guerrero, Mcallister, & Wilson, 2015; Pomeranz et al., 2014).

Over the last decade, the ecological data available on the distribution, abundance, and movements of migratory birds have dramatically increased. There has been a renaissance in our understanding of bird migration due to recent technological advancements. Tracking technologies are now revealing intricate migratory pathways, complex networks of stopover locations, and patterns of migratory connectivity (Knight et al., 2021; McKinnon & Love, 2018; Whitney, 2022). Novel analytical methods include network analyses based on genetic and demographic data (Ruegg, Harrigan, Saracco, Smith, & Taylor, 2020; Xu et al., 2020) and approaches that use seasonal abundance or climate suitability data in spatial prioritizations (e.g., Johnston et al. 2020, Grand et al. 2019). New data products also include the eBird Status maps that provide weekly estimates of relative abundance and occurrence for >1,000 bird species around the globe (Fink et al., 2020) and are a valuable tool for describing migratory bird distributions. The relative abundance maps have been shown to improve prioritization results for conservation planning compared to efforts using only occurrence-based range maps (Johnston et al. 2020) and have been used to develop a variety of hemisphere-scale spatial prioritization approaches (Johnston et al., 2020; Schuster et al., 2019; S. Wilson et al., 2019; Scott Wilson et al., 2022). Exciting opportunities have emerged that integrate these efforts in novel ways. Recently, Meehan et al. (2022) developed an approach to integrate band re-encounter, tracking, migratory connectivity, and eBird Status relative abundance data into a single spatial layer for a given species, revealing new insights into species-specifiec spatial distributions during migratory periods. Each of these innovations advances our ability to prioritize places for conservation actions to benefiet migratory birds.

Although these innovations have contributed to hemisphere-scale conservation planning, application of this information to identify sites for on-the-ground conservation actions remains complex (Martin et al. 2007). The application of this broad-scaled work to local perspectives is essential and Wyborn and Evan (2021) go so far as to question the necessity of more global prioritizations without clear links to localized conservation work. Ultimately, the places where conservation actions occur must be relevant locally, yet informed by processes that occur regionally, hemispherically, or even globally (Cash et al., 2006; Efrat, Hatzofe, & Berger‐Tal, 2020; Santangeli et al., 2020). Local actions (e.g., land protection, ecological restoration, or invasive species management) should be considered in the context of the full annual cycle for migratory birds to ensure that each season is sufficiently addressed when planning conservation actions (Hobson & Wilson, 2020; Marra, Cohen, Loss, Rutter, & Tonra, 2015).

Translating hemisphere-scale priorities into local conservation actions presents an opportunity for co-production with the communities that will ultimately be implementing these actions (Chambers et al., 2021). Conservation efforts that are informed by broad-scale science face challenges if the interest and capacity do not exist for those who may ultimately be implementing the conservation actions (Enquist et al., 2017; Saunders et al., 2021). This issue may be of particular concern for wide-ranging migratory species that span political boundaries and are informed by relatively coarse-resolution prioritizations. Translational ecology, an approach where science producers and science users co-produce research that is tailored specifiecally to meet the needs of the end users, offers a practical framework for centering the needs of resource managers and conservation decision makers to direct science (Enquist et al., 2017; Saunders et al., 2021). Within this framework, conservation practitioners (hereafter, practitioners) who will ultimately use and apply the science-based tools, must meaningfully participate in each major step of the process. Non-governmental organizations present a unique opportunity to implement translational ecology, as it is common for both science producers and users to be part of the same organization (See Fig. 1, Saunders et al. 2021). Translational ecology is also an integral component to the process of applying broad-scale prioritizations to on-the-ground conservation action. Here we describe a multi-level framework for using hemisphere-scale, full annual cycle prioritizations to inform on-the-ground conservation actions. For this application, we defiene ‘multi-level’ simply as being organized into levels, and do not infer a formal statistical approach or particular chronology. Our framework could be applied by any organization(s); however, this work was conceived in the context of a non-governmental organization with a hemispheric reach, the National Audubon Society. The National Audubon Society’s conservation efforts include both broad-scale national and international, as well as local, site-based conservation programs. To ensure that its conservation programs are informed by science, Audubon has invested in spatial analyses across multiple scales (e.g., Audubon Alaska 2019, Grand et al. 2019; Saunders et al. 2019; Bateman et al. 2020b; Michel et al. 2020; DeLuca et al. 2021; Michel et al. 2021). The utility of these investments is measured by the degree to which they can support state and regional efforts throughout the hemisphere to advance on-the-ground conservation and policy. As declines in migratory birds continue (Rosenberg et al. 2019), the National Audubon Society has recognized the importance of connecting hemisphere-scale, full annual cycle perspectives to local conservation actions.

**Figure 1.**
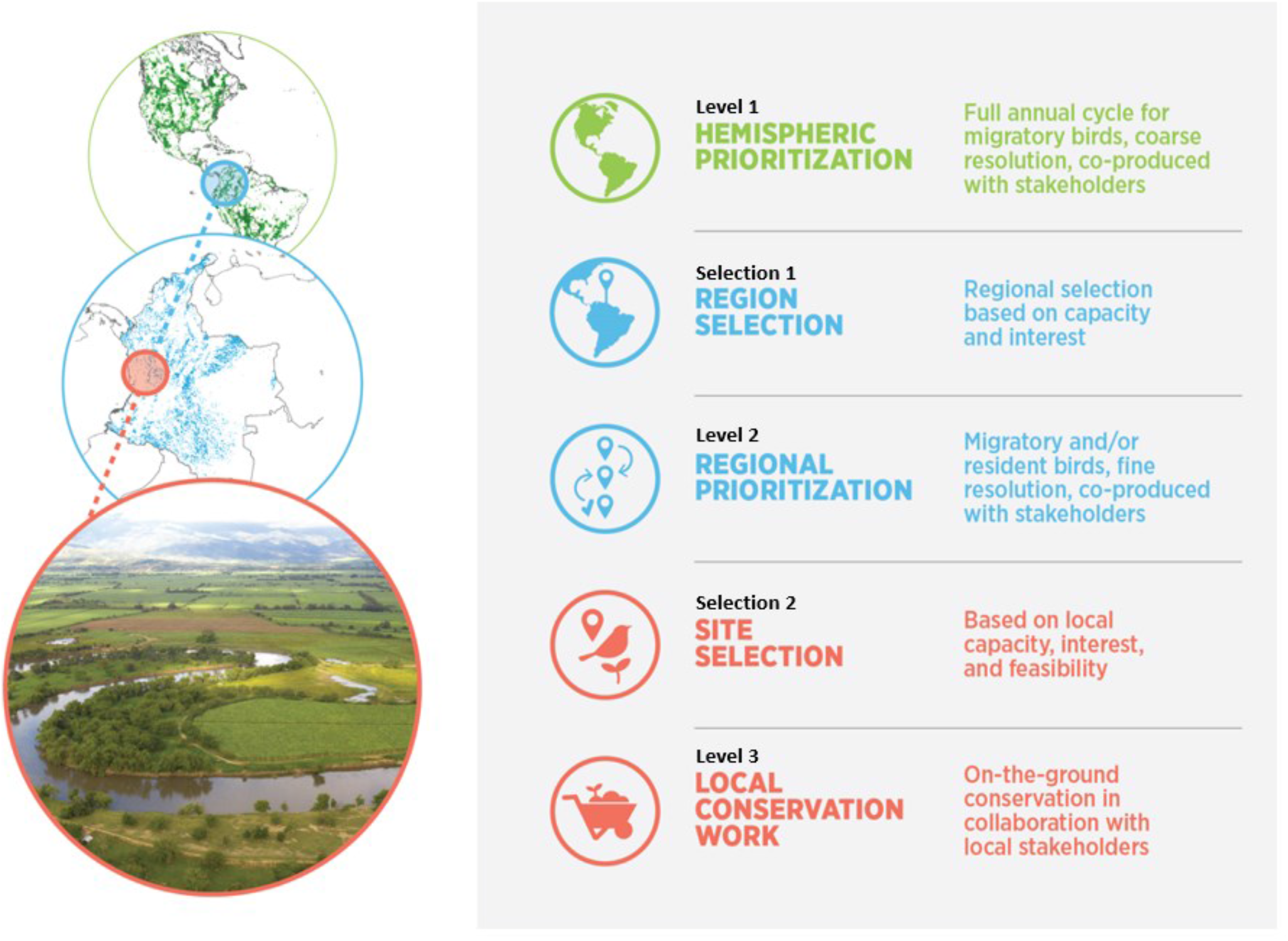
A multi-level framework for linking coarse resolution, hemisphere-scale, full annual cycle prioritizations with regional and ultimately local conservation work. Level 1 - Prioritizations are developed at the hemisphere scale to identify important regions for the full annual cycle of migratory birds. Selection 1 - Organizational capacity and interest as well hemispheric priorities are used to select important regions. Level 2 - Finer resolution, regional prioritizations are conducted and focus on local practitioner interests or conservation targets. Selection 2 - Local scale sites are selected based on organizational capacity and community/partner engagement. Level 3 - Site-based work is conducted in collaboration with local practitioners and organizational partners.

The framework (Fig. 1) includes fieve components, consisting of three levels and two selection steps between each level: Level 1 - develop full annual cycle spatial prioritizations for migratory birds, Selection 1 - select important regions based on broad-scale prioritizations and organizational capacity, Level 2 - develop fiener resolution, regional prioritizations that match local conservation interest, Selection 2 - select local sites to apply relevant conservation actions, Level 3 – local, on-the-ground conservation actions. Throughout this process, we use a translational ecology perspective and highlight how and where practitioner interactions can help ensure that conservation science is actionable. It is not essential that the level 1 and 2 prioritizations are developed in a particular order, rather that the regional work is interpreted within the context of the hemispheric, full annual cycle prioritization. To illustrate the application of the framework, we describe two examples from real-world conservation planning effort. One example is from the coastal region of North and South Carolina, USA, and the other for Colombia.

## APPROACH

### Western Hemisphere

#### Level 1 - full annual cycle prioritization for North American breeding migratory birds

To incorporate full annual cycle information into Audubon’s conservation decision making processes for migratory birds, we fierst identifieed six major focal ecosystems for birds that breed in North America: boreal forests, western forests, arid lands, grasslands, eastern forests, and wetlands/coasts. These ecosystems represent distinct habitat types and corresponding bird communities. These ecosystems also align with Audubon’s existing conservation initiatives. For each of these ecosystems, we created a list of target species with the number of species ranging from 21 (boreal) to 91 (wetlands/coasts; Table S1). We selected species based on the following criteria: within an ecosystem, species should represent the range of habitats and conditions that meet the nesting and foraging needs for most breeding birds in that ecosystem; species must be migratory; and species should be relevant to conservation work at Audubon and other conservation entities where possible (e.g., US Fish and Wildlife Service, Partners in Flight, etc.). Some species that breed in more than one habitat type were selected for multiple ecosystems. Ecosystems were used solely to inform the species selections and did not limit the geographic extent of the prioritizations. Once species were selected for an ecosystem, their full range throughout the annual cycle was considered. For example, Blackpoll Warbler (*Setophaga striata*) was selected to represent both eastern forests and boreal forests groups; therefore, the full breeding, nonbreeding, and migratory ranges were included for both prioritizations.

We conducted a separate spatial prioritization for each ecosystem (Fig. 2) using Zonation conservation planning software (Moilanen et al., 2014). Zonation provides several heuristic optimization algorithms that rank the conservation value of cells in a landscape using systematic conservation planning principles such as complementarity, representation, and efficiency. We used Zonation for several reasons; fierst, Zonation does not require the user to set population targets, rather it employs an iterative removal process beginning with the least valuable cells. This approach more closely aligned with our study objectives. Second, Zonation allowed us to intuitively incorporate subregions to ensure a spatially balanced network of priority areas that were of interest to the practitioners. Finally, Zonation is a spatial prioritization software that is both accessible and familiar to most practitioners. We used the Core Area Zonation ranking method, which ranks cells by iteratively removing the cell with the lowest marginal value across the landscape. Cells with the highest quality for any one species, as well as those that simultaneously benefiet many species, receive the highest ranks. The result is a landscape ranking that ranges from 0 to 1 (Moilanen et al., 2014). Administrative units, primarily based on Bird Conservation Regions in North America and Ecoregions in Central and South America (Figure S1; Meehan et al. 2022), were used to stratify the analysis ensuring that the full range of landscape ranks were well distributed across the hemisphere. This is particularly important for migratory birds because a network of suitable habitat that spans their full annual cycle is essential. However, because the full gradient of values (0-1) is ensured for each unit, spatial comparisons at the country scale for example, have little relevance. All administrative units and all species were weighted equally in Zonation. For each species, four separate spatial data layers were used as inputs to Zonation. For the breeding and stationary nonbreeding seasons, we used seasonal mean relative abundance (i.e., the mean pixel-level relative abundance averaged across the weekly layers for each season) from eBird Status data (Fink et al., 2020). Although eBird abundance models provide remarkable distribution and abundance information across species’ ranges, in remote regions (e.g., some portions of the boreal forest and the Amazon Basin), inadequate survey and remotely sensed data prohibit accurate abundance predictions (Fink et al., 2020). Any error present in the eBird abundance model inevitably propagated to our prioritizations. We used separate layers for spring and fall migration that integrate tracking, banding, migratory connectivity, and eBird Status relative abundance data (Meehan et al. 2022). This integration process used probabilistic least-cost paths and a generalized additive mixed modeling approach to describe spatial patterns of avian migration and resulted in merged migration maps with a resolution of 27 km (Meehan et al. 2022, Figs. 2 and 3 for an example). Each pixel was then assigned a value (0-1) representing an index of concentration during the migratory periods. Importantly, each seasonal spatial layer for each species was treated as a separate feature in the prioritization, ensuring that the most important places during each season, across the hemisphere, were assigned the highest ranks. The resolution of the resulting prioritizations was 27 km due to the resolution of the merged migration maps (Meehan et al. 2022). To facilitate interpretation of the relatively coarse data and smooth artifiecial boundary effects between pixels and administrative units, we used bilinear interpolation to downscale the data to 3 km resolution and then applied a Gaussian-weighted kernel smoothing algorithm with a 25-km standard deviation. We used the library ‘gridkernel’ in Program R v3.6.1 (R Core Team, 2020) to conduct the spatial smoothing. It is important to note that although the resulting raster layers have a cell size of 3 km, the functional resolution remained at 27 km and we interpret these results at landscape or regional scales, on the order of >100 km2. Once each of the six ecosystem prioritizations (Fig. 2 a-f) were processed in this manner, we calculated the maximum value at each cell across the six layers. For interpretation and to facilitate regional selection (Fig. 1, Selection 1) for further consideration, we then mapped the 90th, 80th, and 70th percentiles (Fig. 3).

**Figure 2.**
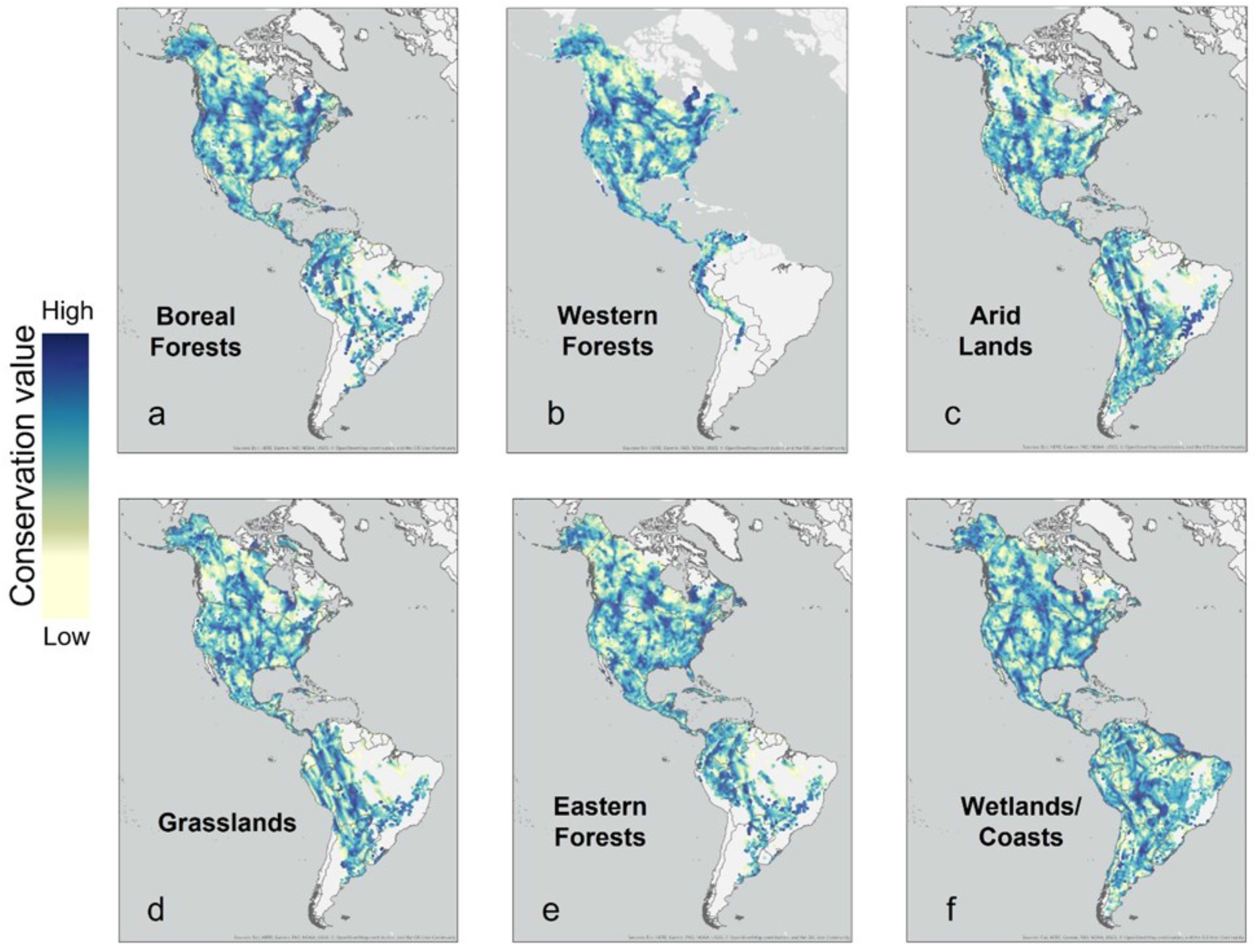
Full annual cycle spatial prioritizations for six separate breeding ecosystems, totaling 171 species, representing (a) boreal forests, (b) western forests, (c) arid lands, (d) grasslands, (e) eastern forests, and (f) wetlands/coasts. Prioritizations for each ecosystem group (a-f) were a function of the target species; no spatial limitations were applied. For example, Blackpoll Warbler was selected for the eastern and boreal forest prioritizations and was therefore mapped across its range spanning the boreal forest across North America and throughout the hemisphere to capture its full annual cycle. Prioritizations were performed using the conservation planning software Zonation and the results are illustrated on a scale from 0 to 1 representing low to high conservation value. White spaces indicate no data for that prioritization. See Table S1 for the species list used for each map.

**Figure 3.**
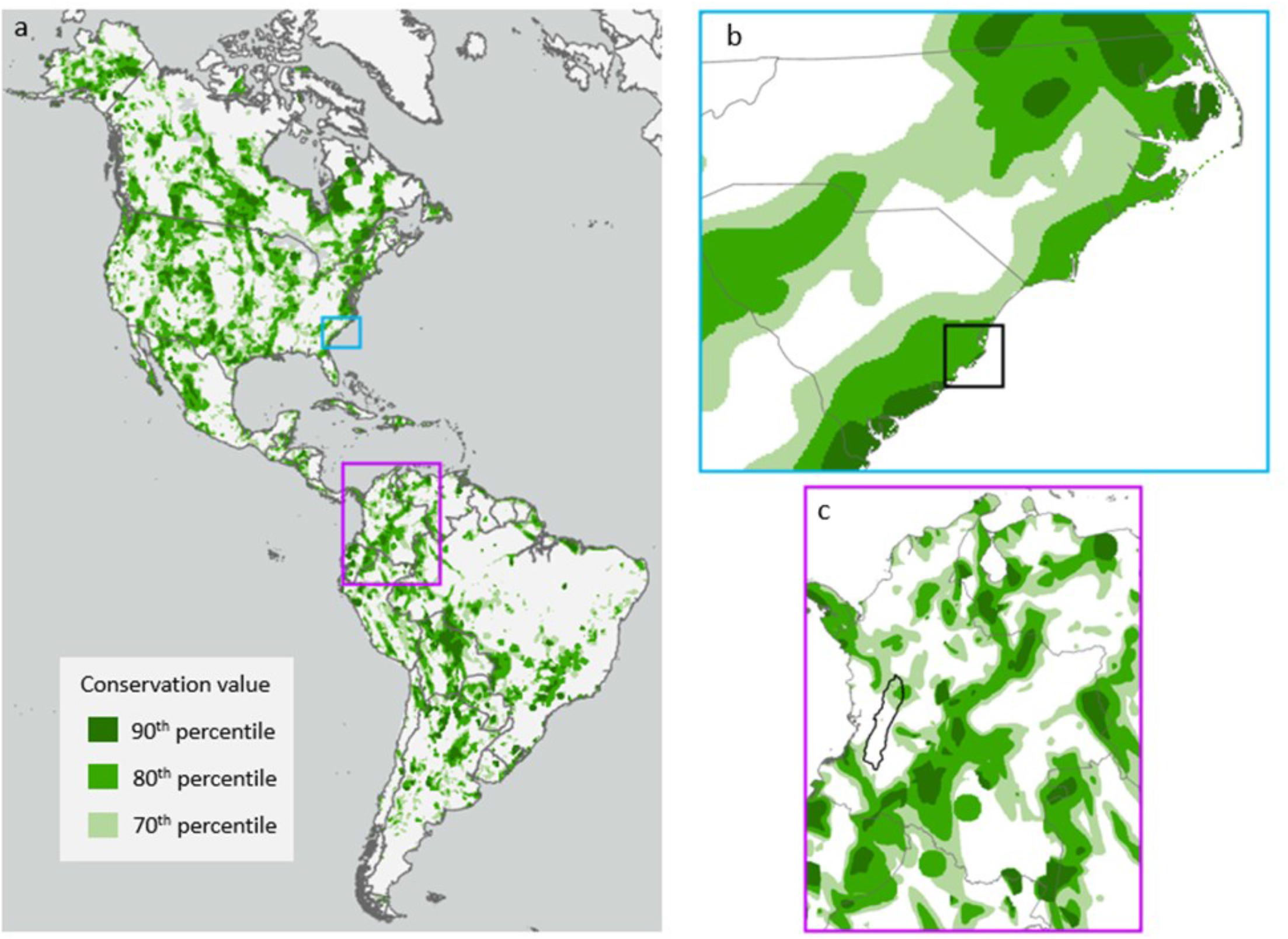
(a) The maximum prioritization rank value across six full annual cycle prioritizations (Fig. 2), summarized for the 90th, 80th, and 70th percentiles. This fully integrated prioritization was then used to identify important regions for finer resolution analysis. The Coastal Carolinas (b) and Colombia (c) were then selected for finer resolution prioritizations. Highlighted in black are the locations selected with the separate regional prioritizations. See Figures 4 and 5.

Practitioners provided valuable contributions at several key decision points in the process of developing the full annual cycle prioritizations. At this stage (i.e., Level 1 and Selection 1; Fig. 1), practitioners consisted of Audubon staff with diverse expertise and geographic focus. They were consulted monthly across an eight-month period and informed the following decisions: (1) The selection of the ecosystems that ultimately determined the species lists. Ecosystems were selected to best align with and inform regional initiatives. (2) The selection of species used in each ecosystem prioritization based on the general criteria described above. Practitioners carefully considered the relevance of each species to their work and to the interests of the local communities they engage with. (3) The decision to weight each species equally within Zonation. (4) The downscaling and smoothing of the 27 km resolution raster layers to 3 km to improve user interpretation of the spatial prioritizations.

The analysis resulted in priority ranks across the Western Hemisphere for all locations where the 172 species included in the prioritizations were predicted to occur throughout their full annual cycles. The prioritization approach we selected balances the value of places that are important to multiple species and those that are important to single species or seasons that are not included in areas of high species richness (Moilanen et al., 2014). The top-ranked regions highlighted by the full annual cycle prioritizations comprise a representative and efficient network of sites that capture important locations for all species during all seasons of the year in a minimal area and can be considered for further prioritization in a regional context.

The integrated full annual cycle prioritization (Fig. 3a) was generally consistent with our expectations of the important places for migratory birds. Some examples include the Prairie Pothole and Great Salt Lake regions, which were both highlighted due to their importance to the waterbird communities; the northern mid-Atlantic New England and Quebec regions, due to their importance to the breeding eastern forest and migrating boreal forest bird communities; Hispaniola for migrating and overwintering water, boreal and eastern forest birds; and central Bolivia for overwintering forest, grassland, arid land and waterbirds. There are also some notable regions that were not identifieed by the full annual cycle prioritization. For example, the Amazon Basin was not identifieed because none of the species included in the prioritizations (Table S1) are dependent on the Basin during their annual cycle. In fact, the Amazon Basin generally has low abundance of Neotropical migratory birds (Robinson, Fitzpatrick, & Terborgh, 1995).

#### Selection 1 - Connecting the hemisphere and the region—Coastal Carolinas and Colombia

In our framework (Fig. 1), the full annual cycle prioritization can be used to identify regional locations of importance for further consideration (Fig. 3). When interpreting the full annual cycle prioritizations, it is important to note that they are intended to identify broad regions for further conservation consideration. For this purpose, we consider regions to be on the order of tens of millions of hectares (state or country scales rather than towns or parcels) and relevant to broader conservation initiatives. It follows that practitioners would be most interested in regions identifieed by hemisphere-scale prioritizations that overlap with organizational conservation capacity and goals. Here we outline two examples of this process (see also Wilson et al. 2022 as an example of this phase of the framework) in the Coastal Carolinas, USA (Fig. 3b) and Colombia (Fig 3c). These two geographies emerged as areas for greater investment for three reasons. First, the full annual cycle prioritizations identifieed them as important landscapes for migratory birds (Fig. 3). Here we use the 70th, 80th, and 90th percentiles (as described above) to highlight two important regions. The Coastal Carolinas (Fig. 3b) are almost completely encompassed within the 70-90th percentiles and a large portion of Colombia was also within the 70th – 90th percentiles (Fig 3c). Second, there was organizational capacity to develop and apply regional, fiene-scale resolution analyses for these locations. Although many areas were identifieed across the hemisphere as being important, one organization simply does not have the capacity to work on them all. Therefore, considering realistic organizational capacity is important. Third, there was interest, from local practitioners, in identifying sites for local conservation action. Without local interest in on-the-ground conservation efforts, desired conservation outcomes are unlikely.

### Coastal Carolinas

#### Level 2 – Regional prioritizations

The Coastal Carolinas region was highlighted within the 70th - 90th percentiles of the full annual cycle prioritization (Fig. 3b). In this region, Audubon North and South Carolina were interested in developing a regional prioritization that spanned across the two states. The regional prioritization was co-developed by Audubon South Carolina, Audubon North Carolina, and Audubon’s national science team. The goal of the Coastal Carolina’s spatial analysis, as defiened by practitioners that included staff from both the Audubon North and South Carolina fieeld offices and local Audubon chapters, was to guide strategic conservation planning across the Coastal Carolinas. They desired to focus on: 1) importance for priority birds accounting for climate change, 2) ecological integrity, and 3) co-benefiets for local human communities.

For this regional prioritization, we created two species lists, one for marsh birds and one for coastal birds (seabirds and shorebirds; see Table S2). Separately for each guild, we created present and future (mid-century) prioritizations using Zonation conservation planning software (Moilanen et al., 2014). The geographic extent of each prioritization was defiened by NO-AA’s marsh migration model, which delineated locations of present and predicted future marsh and coastal habitats (Office for Coastal Management, 2020). Target features included present and future bird distributions, delineated using species distribution models developed by the National Audubon Society (1 km resolution) that accounted for both climate (RCP 8.5) and habitat suitability (Bateman, Taylor, et al., 2020; Wilsey et al., 2019). We also included nest point locations obtained from censuses conducted by Audubon North Carolina and the South Carolina Department of Natural Resources during 2000-2020. Additional present and future target features included resilient marshes and marsh migration areas, respectively, (Anderson & Barnett, 2017) and the distribution of subaquatic vegetation compiled by the North Carolina Department of Environmental Quality. We weighted all species equally and weighted avian target features twice that of non-avian targets. We combined the global human modifiecation index (Kennedy, Oakleaf, Theobald, Baruch‐Mordo, & Kiesecker, 2019) with recent surface water increase (Pekel, Cottam, Gorelick, & Belward, 2016) to create a present landscape condition layer, and combined predicted urbanization (U.S. Environmental Protection Agency, 2017) and flooding threats (Chesnutt, Dobson, Johnson, Rhodes, & Hutchins, 2019) to create a future landscape condition layer.

To identify areas where habitat improvements could benefiet both birds and local coastal communities, we used the Center for Disease Control’s Social Vulnerability Index (Flanagan, Gregory, Hallisey, Heitgerd, & Lewis, 2011) as a cost layer such that census tracts with high social vulnerability were considered low cost and thus ranked higher (cost = 1-SVI). We used the Core Area Zonation algorithm to rank the present and future landscapes (0-1), based on important sites (core areas) for all focal birds, that have high ecological integrity, and are in close proximity to socially vulnerable communities. Practitioners centered the social vulnerability component because vulnerable coastal communities often face increased exposure to climate change hazards and decreased capacity to recover from those hazards, which necessitates that conservation planners and practitioners balance human and ecological needs in return for increased benefiets to both.

To identify suitable management actions for priority locations, we combined the results from the present and future prioritizations into a management classifiecation layer (Fig 4). Locations ranked in the top 30% in both present and future time periods were classifieed as top priorities for protection or maintenance; locations ranked in the top 30% in the present but not the future were classifieed as top priorities for adaptation; locations ranked in the top 30% in the future but not the present were classifieed as top priorities for restoration; and locations ranked in the bottom 70% in both time periods were classifieed as low priority. This four-category management classifiecation, in combination with local knowledge, is currently being applied to guide on-the-ground conservation efforts in the Coastal Carolinas region.

**Figure 4.**
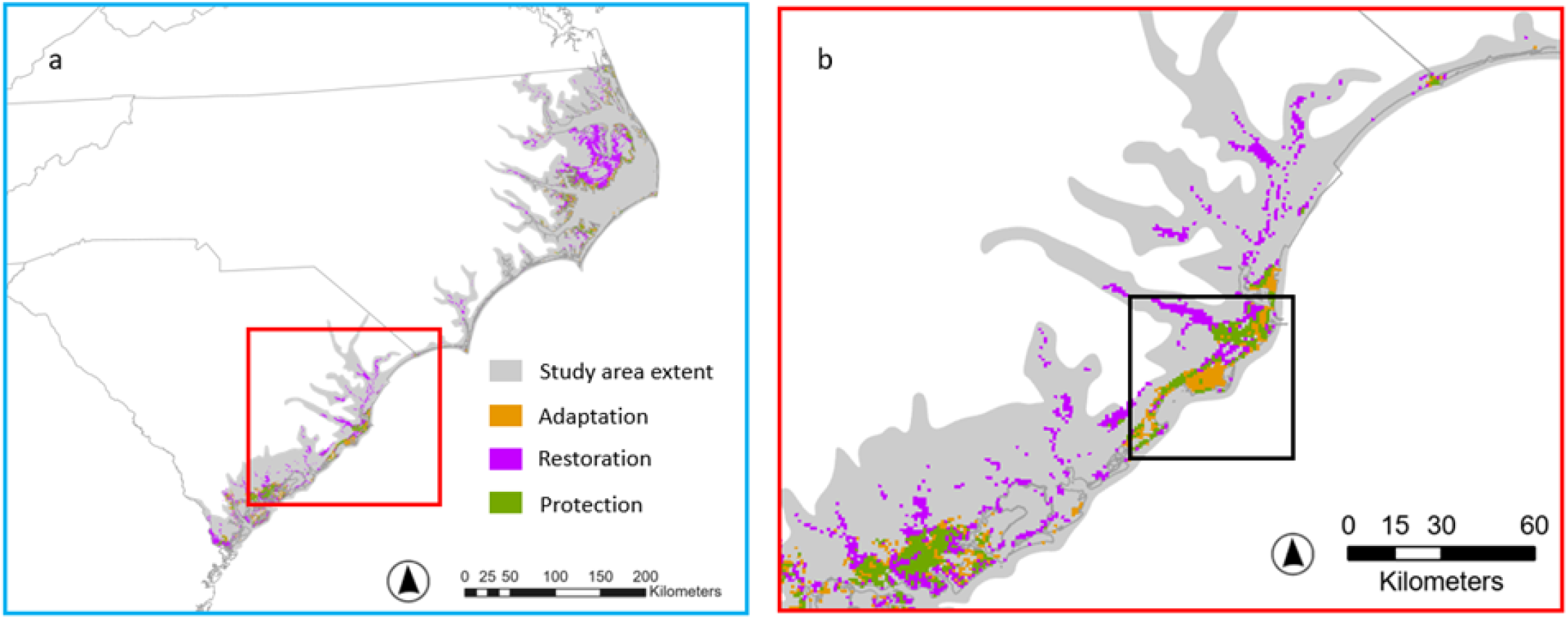
(a) The results of a localized, fine resolution, prioritization to identify areas along the coast of North Carolina and South Carolina, USA for either habitat adaptation, restoration, or protection for migratory shorebirds. This prioritization was conducted using the conservation planning software Zonation. The selection of the extent of this prioritization could be informed by a broader, hemisphere-scale prioritization, as shown in Fig. 3. (b) The black box shows the location for planned community engagement and marsh resiliency work.

#### Selection 2 and Level 3 – From regional prioritizations to local work

The Cape Romain National Wildlife Refuge and surrounding area (Fig. 4a) has a high concentration of locations ranked in the top 30% of the regional Coastal Carolina prioritization as indicated by areas recommended for adaptation, restoration, and protection for beach and marsh habitat compared to other areas of the coast (Fig. 4). In addition, Audubon’s South Carolina fieeld office has existing capacity (resources, community networks, and institutional infrastructure) near the Cape Romain National Wildlife Refuge; thus, this area was selected to focus limited conservation resources. Preliminary work will focus on coastal resiliency and community engagement efforts in the Cape Romain area, which may include natural infrastructure projects to prevent marsh erosion, facilitate marsh migration, and slow the loss of island nesting sites to maintain and improve habitat for breeding and migrating marsh and shore-birds. The regional prioritization has also helped Audubon South Carolina build relationships with a cadre of partners because it has served as a visual representation of a strategic plan for resilience work, allowing local work to focus on the places that balance social and political opportunities with the priority ranks to target the most important places for birds and people. Audubon South Carolina is working to engage such partners within the community as the Army Corps of Engineers, private engineering fierms, the South Carolina Ports Authority, the US Fish and Wildlife Service, the South Carolina Office of Resilience, the South Carolina Department of Natural Resources, and other conservation NGOs. The co-creation of coastal resilience projects with these community partners will ensure that multiple practitioners are invested in the success of the conservation projects.

### Colombia

#### Level 2 - Regional prioritizations

Colombia was identifieed as a priority in the full annual cycle prioritization for wetland, eastern forest, boreal forest, and grassland birds (Fig. 2 and 3). In Colombia, Audubon Americas (AA) was interested in developing a fiener resolution prioritization that could guide conservation efforts to benefiet both migratory and resident birds. As part of the development of AA’s business plan, a spatial prioritization was developed to select “no-regrets” geographies (called “deep intervention landscapes”) for its Working Lands strategy in Mexico, Panama, Colombia, and Chile. These geographies are designed to highlight areas where AA and its partners can demonstrate bird-friendly and sustainable agricultural practices. AA began with a list of 119 North American migratory species that had been prioritized by Audubon as well as 303 declining migratory species (sensu Rosenberg et al. 2019). From this set, 108 species had over 50% of their population at any point of their annual cycle within AA countries, as determined by comparing the summed relative abundance from eBird status layers (Fink et al. 2020) across the species’ range to the relative abundance within AA countries for each season (nonbreeding, prebreeding and postbreeding migration). We also considered in the prioritization, resident species that were either threatened according to BirdLife International (2020) or country endemics, based on national checklists (Avendaño et al., 2017; Berlanga et al., 2015) and Avibase (https://avibase.bsc-eoc.org/). In Colombia, these criteria resulted in 42 migratory species and 178 resident species that were either threatened, according to BirdLife International (2020), or country-level endemics based on national checklists (Avendaño et al. 2017). For migrant birds, we gathered distributional data from eBird status models (Fink et al., 2020) whereas for resident birds we developed area of habitat models based on the workflow proposed by Palacio et al. (2021). Representation targets for each species were set at 10% population for migrant landbirds, 30% for migrant waterbirds, and a sliding target between 10-100% of the distributional range of resident birds based on their current range sizes. The population thresholds are within the bounds of previous work (Johnston et al., 2020; Rodrigues et al., 2004; Schuster et al., 2019) and were agreed upon by practitioners. We used Gurobi software through the R package prioritizr (Hanson et al. 2020) to identify the minimum set of areas (3 km cells) that would achieve the representation targets of all species considered across Colombia. We did not consider cost of conservation actions in each cell, as practitioners were interested in identifying the most important areas for birds unconstrained by cost. It is important to highlight that this prioritization approach differs from the approach used in Level 1 and was designed specifiecally to meet the objectives of the practitioners at this regional scale.

The approach above yielded a binary indicator of cells being part of a solution that minimizes area to meet targets (i.e., 1) or not (i.e., 0). We aggregated this solution into landscapes (using watersheds from HydroBASINS level 6 as our operational unit; Lehner & Grill 2013) and computed their contribution to meeting targets, as well as current levels of threat using the average human footprint map (Venter et al. 2018) of the cells identifieed as part of the solution within each landscape. We prioritized conservation action for landscapes with high levels of threat and contribution to meeting targets.

#### Selection 2 & Level 3 - From regional prioritizations to local work

The regional prioritization for Colombia’s migrant and resident bird communities resulted in 12 prioritized landscapes in Colombia and of those, two priority landscapes were selected for localized conservation actions. Here, we focus on the Cauca Valley landscape (Fig. 5a). Although the entire Cauca Valley was not highlighted in Level 1, it was selected based on the prioritization conducted for Level 2 as well as local practitioner interest and capacity to work in the Cauca Valley as a regional conservation planning unit. The fiener resolution prioritization (Level 2) coupled with practitioner input led to the selection of a focal region (Cauca Valley), that the hemisphere-scaled prioritization alone could not have identifieed. Additionally, selecting these landscapes was meant to ensure their contribution to meeting national bird conservation targets. Changes in species’ distributions and abundances elsewhere in the country (e.g. due to landcover changes) that may occur during implementation in the selected areas are not considered in this approach, which is typical for minimum set coverage conservation problems (Meir, Andelman, & Possingham, 2004; Possingham, Moilanen, & Wilson, 2009). For local conservation actions in the Cauca Valley watershed, we ranked sub-watersheds (HydroBASINS Level 7; Lehner and Grill 2013) according to their contribution to meeting nationwide targets, and feasibility which was assessed via internal consultation to identify areas with established community associations with which AA could partner. In selecting this focal area, we aimed to maximize contributions to bird conservation while using existing governance and fienancial tools to accelerate the adoption of bird-friendly practices in working lands and protection of existing natural habitats. Within the Cauca Valley, AA worked with local practitioners to select two sub-watersheds to begin work (Figure 5b). The variation in the conservation value of these sub-watersheds illustrate the complexities of applying prioritizations to on-the-ground decision-making. The sub-watershed selected in the northern section of the valley was clearly highlighted by both the hemispheric and regional prioritizations (Fig. 5b). However, the sub-watershed selected in the south was not highlighted by either prioritization. This sub-watershed was selected because practitioners identifieed it is a critical water source for the area to the southeast, which was highlighted by the prioritizations. Additionally, practitioners in the region have existing relations with local communities in the sub-watershed, which contributed substantial capacity and ultimately lead to the selection of this sub-watershed for further conservation work. In these two sub-watersheds, practitioners will identify farms that are interested in transitioning to bird-friendly practices without compromising farm production. Such practices may include planting hedgerows with edible plants for cattle, planting fruiting trees for birds, using silviculture to increase farmer’s productivity while halting expansion of pastures to forest remnants, planting live fences with trees useful for birds, and fencing forest remnants to avoid cattle degradation of the understory (Figure 6). Based on the lessons learned through the implementation of conservation efforts in these sub-watersheds, future conservation work can be directed to watersheds that are both hemispheric and regional priorities.

**Figure 5.**
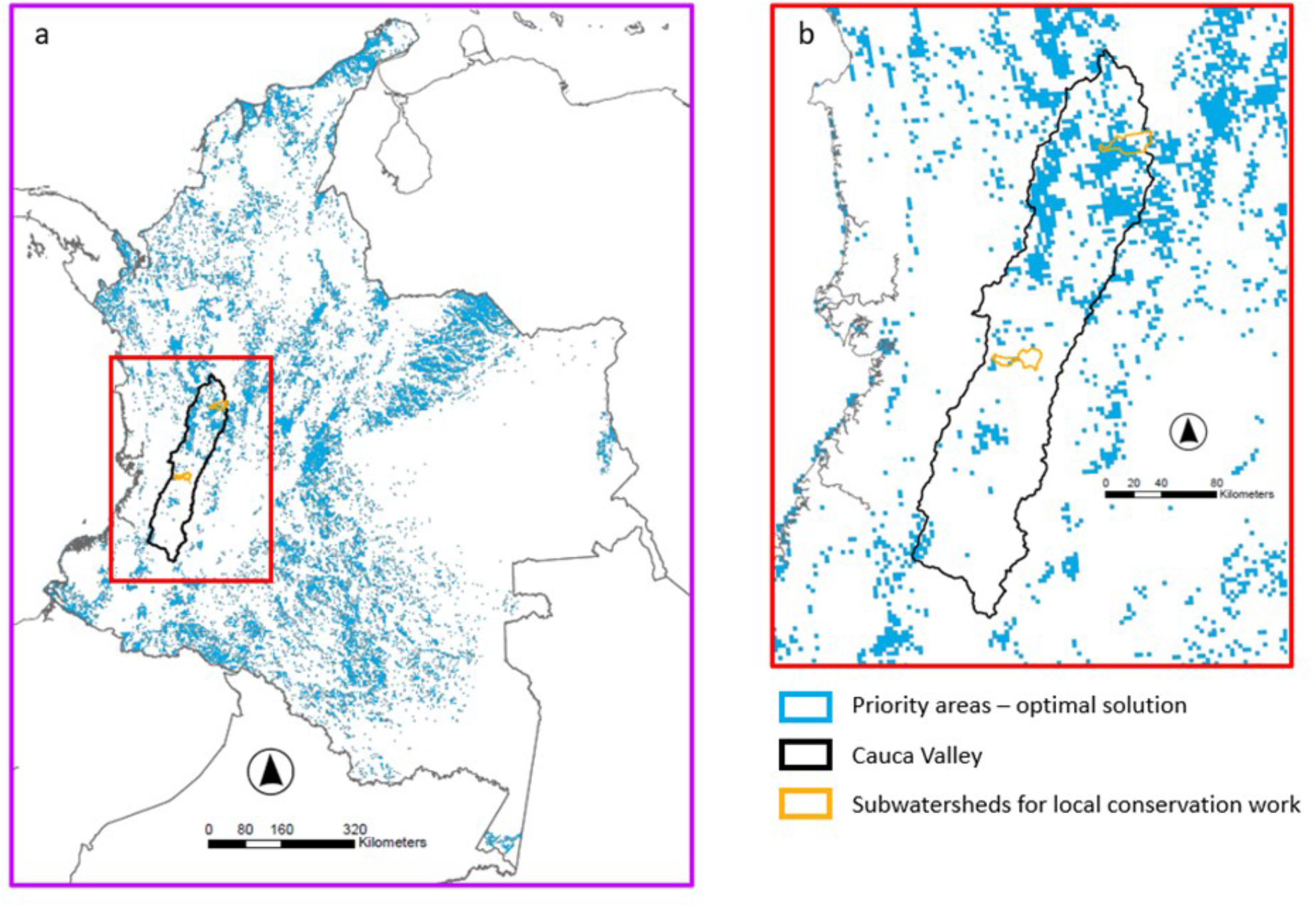
(a) The results of a localized, fine resolution, prioritization to identify important conservation areas using 175 resident and 42 migratory bird species in Colombia, South America. The selection of the extent of this prioritization could be informed by a broader, hemisphere-scale prioritization, shown in Fig. 3. The prioritization for Colombia was developed to meet specific population targets and implemented in the R package ‘prioritizR’. Values represent the frequency cells were selected in near optimal solutions, thus representing conservation value. (b) Based on the prioritization and conservation practitioner input, the Cauca Valley was selected for further consideration. Specifically, two subwatersheds were selected for localized conservation work.

**Figure 6.**
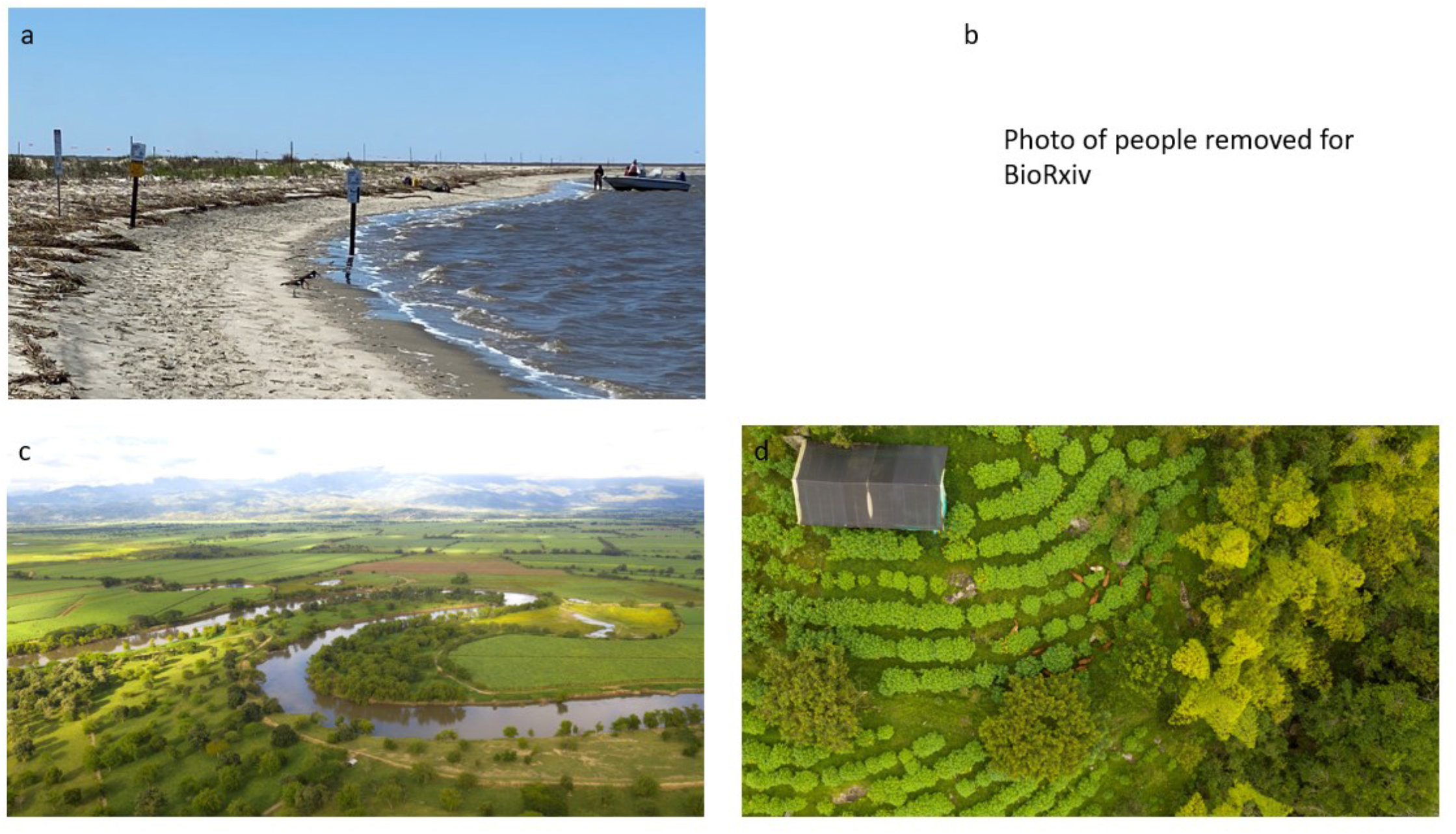
Photographs where local conservation work is occurring based on regional prioritizations (Fig. 5 and 6). a and b show local stake-holders cooperating to post information on beach nesting birds in and around Cape Romain National Wildlife Refuge. (c) Cauca Valley, Colombia where local conservation work has begun to integrate farming and bird habitat conservation. (d) Site where farming and bird habitat conservation coexists.

## DISCUSSION

The multi-level framework described here for migratory birds (Fig. 1) provides a roadmap for considering full annual cycle processes at hemisphere scales that can guide regional prioritizations that lead to local conservation actions. Conservation activities ultimately impact wildlife and people in specifiec places (Pressey et al. 2013); thus, translating broad-scale conservation plans into the local context, while maintaining practitioner engagement, is critical for ensuring that conservation efforts achieve the desired outcomes (Winter et al. 2006; Morissette et al. 2019) and that these activities do not have negative impacts or unintended consequences for local communities (Holmes, 2013; MacKenzie et al., 2017). Our framework explicitly integrates feedback from practitioners during each step, ensuring that science users are invested in the process. This framework aims to accommodate real-world, practitioner-driven decisions that account for institutional capacity and local needs, while ensuring that the basis for those decisions is grounded in science.

Our framework contributes to a growing set of approaches to address the challenge of incorporating multiple scales of information into spatial conservation planning (Cash et al. 2006). There have been two general approaches to this challenge, one has been to incorporate data from multiple spatial scales and resolutions into a single prioritization (Lagabrielle et al. 2018, Rinnan et al 2020). The single prioritization approach is most advantageous when quantitatively linking information from each spatial scale is critical and when practitioners’ needs are aligned across scales. The other general approach, which we have illustrated here, has been to conduct separate analyses at increasingly fiener spatial resolutions (e.g., Hernández-Matías et al. 2020). This approach can be used to guide conservation efforts on single species, for examples Hernandez-Matias et al. (2020) prioritized areas to take actions to reduce Bonelli’s Eagle (Aquila fasciata) mortalities by beginning with a metapopulation analysis at the broadest spatial scale, and then focusing a separate analysis at the territory scale and ultimately identifying core areas. While not a spatial prioritization, per se, this approach highlights the flexibility of incorporating distinct analyses at each scale to address a specifiec conservation challenge. Often, in migratory bird conservation, the differences between the hemisphere and local scales in terms of spatial resolution, ecological and social spatial data layers, practitioner needs, conservation challenges, and expected outcomes, are so vast that maintaining flexibility at each step of the process is essential. Our multi-level framework achieves this flexibility by incorporating tailored practitioner contributions at each level, while also enabling science providers to implement the most appropriate analyses to best address the practitioner requests.

The nature of our framework (Fig. 1) allows practitioner groups to vary at each level of the process and from region to region. We built hemisphere-scale prioritizations with practitioners interested in using them to identify broad regions that were important for migratory bird conservation to direct regional strategic plans, reinforce existing projects, and identify new conservation initiatives. Due to the broad geographic extents, direct connections to local conservation actions and the affected communities can be unrealistic at hemisphere scales. Furthermore, local scale practitioners are likely to be nearly entirely different from one region to the next. The decisions made at Level 1 and Selection 1 of the framework (Fig 3) ultimately serve to focus conservation efforts at regional scales, which often still span political and organizational boundaries but are more in line with developing regional strategic conservation plans. Therefore, it is important to engage different or increasingly diverse practitioners to conduct and apply the regional prioritizations (Young et al., 2013).

At regional and local scales, Levels 2 and 3 of our framework (Fig. 1), practitioners are more connected to the people, politics, and places where on-the ground conservation actions occur. Furthermore, they may have existing relationships with partners that are needed for local coordination and community interactions, which was the case with the Cape Romain example. Importantly, the regional scale is where socially vulnerable communities can be centered in spatial prioritizations that focus on co-benefiets of conservation actions for people at highest risk of adverse impacts from sea level rise, urbanization, and other threats. Once candidate sites have been identifieed from a regional prioritization, the fienal site selection process will require additional outreach and relationship building in these frontline communities to develop new partnerships and gauge local support for implementing conservation or restoration actions. Part of this outreach should include the reference back to the hemisphere-scale prioritizations to help local practitioners connect with the importance of their work in a broader spatial context.

In addition to providing the flexibility to include different practitioners at each level, our framework, linking separate prioritizations, also provided a means by which different conservation targets could be included at each scale. Just as the local practitioners engaged will vary across regions, the conservation goals of those practitioners are also likely to be diverse enough to require that different data and prioritization methods are used at Level 1 and Level 2 of the framework (Fig. 1). The primary driver of our hemisphere-scale prioritization was the full annual cycle conservation of migratory birds, relying on eBird Status relative abundance models, tracking data, and band re-encounter data, with the intent of identifying important regions within a hemispheric context. However, a wide range of possible spatial prioritization approaches, input data, and goals allow for great flexibility in developing these analyses. The two regional prioritizations described here were vastly different, as they were driven by local conservation needs and capacity. The Coastal Carolinas prioritization used the same software (Zonation) as the full annual cycle prioritization, but the input layers, scale of application, and desired outcomes varied between the two. The prioritization developed for Colombia added resident bird data that were not included in either the hemisphere-scale prioritization or the Coastal Carolinas and relied on a different prioritization software to better match the needs of that practitioner group to meet bird population targets. These differences illustrate how each prioritization level can be tailored to the direct needs of their practitioners while simultaneously contributing to the broader framework, so long as the local conservation actions are relevant to the goals at both scales. Where the hemispheric prioritization validates the importance of a regionally-identifieed site, knowing which species are driving the high value, and in what season, can help guide conservation for migratory birds alongside other regional targets.

Although our framework (Fig. 1) was primarily developed and applied within a large non-profiet organization, the approach could be applied across multiple organizations. Broad-scale prioritizations developed by one institution could help identify priorities for regional conservation entities such as Joint Ventures or USGS Climate Adaptation Science Centers (Bisbal 2019), which could then conduct fiener-resolution analyses. These entities are well-suited to apply regional considerations to local scales given their diverse practitioner networks. The flexibility in the analytical approaches, the opportunity to facilitate inter- and intra-organizational collaborations, and the co-development with multiple practitioner groups, highlight the potential effectiveness of the framework for migratory bird conservation applications.

Our hemisphere-scale prioritization for migratory birds builds upon a growing body of work to incorporate new information into conservation plans for migratory birds (Runge et al. 2016, Schuster et al. 2019) by including spatial information from spring and fall migrations that is representative of tracking, banding, and distributional datasets (Meehan et al. 2022). The approach also includes migratory connectivity information, the connections between breeding and wintering populations (Webster, Marra, Greenberg, & Marra, 2005), which is recognized as an important element of conservation planning for migratory species (Dunn et al., 2019; Marra, Hunter, & Perrault, 2011; Martin et al., 2007). Our framework could be extended further to include the strength of migratory connectivity between priority regions to explicitly connect priority regions based on the movements of migratory birds. Integrated population models (Zhao et al. 2020), which synthesize data sources on population demographics (e.g., census, productivity, mark-recapture) to estimate and project population dynamics, and shed light on the factors that may limit populations, can also be useful for strategically informing migratory bird conservation plans across broad spatial scales (Saunders, Cuthbert, & Zipkin, 2018; Zipkin & Saunders, 2018). Moreover, developing methods that account for migration stopover duration is essential to further refiene full annual cycle priorities and exciting advances are being made toward being able to incorporate this information (e.g., Nicol et al., 2023). Future research should continue to explore ways to incorporate tracking information, migratory connectivity, demographic information, and movement ecology into broad-scale conservation efforts. Our work builds upon the established concept of coarse fielter/ fiene fielter approach to conservation planning (Lemelin & Darveau, 2006; Noss, 1987). In our approach, the coarse fielter we developed addresses the movement of migratory birds across the hemisphere and recognizes the importance, as have others (e.g., Schuster et al., 2019), of including this information in conservation planning. We can use this coarse fielter to identify broad regions for further consideration and to place existing regional priorities in the hemispheric context. For the fiene fielter, the prioritization approaches we used included fiener resolution information, regionally important conservation targets, and the input from local practitioners. An alternative application of the coarse and fiene fielter concept of our framework could include the use of the full annual cycle prioritization to reduce a broad set of previously selected locations (e.g., Important Bird Areas or Key Biodiversity Areas) to a smaller subset of sites with the highest priority for migratory bird conservation. However, this approach could miss opportunities to consider local conservation targets and concerns, exemplifieed by the selection of the southern Cauca Valley sub-watershed. In general, our framework offers an adaptable model for integrating coarse fielter and fiene fielter approaches to guide conservation actions on the ground and communicate the contribution that local efforts make to hemispheric challenges.

Across their full annual cycle, North American migratory birds can range from Alaska to Argentina. Across this broad landscape, conservation efforts need to be implemented within the context of local ecology, governance, social values, and conservation threats (Villero, Pla, Camps, Ruiz-Olmo, & Brotons, 2017). However, local efforts must be informed by life cycle processes that span the hemisphere. (Anderson et al., 2018). The approach presented here illustrates how multiscale conservation planning can bring conservation practitioners together to develop locally relevant conservation plans that are nested within hemispheric perspectives. Such multi-level collaborative approaches are urgently needed if we are to successfully reverse the ongoing population declines of migratory birds.

[Data will be uploaded to an appropriate repository upon manuscript acceptance.]

## Supporting information

Figure S1. Migratory Connectivity Regions map.

Table S1. Species list for full annual cycle prioritizaions.

Table S2. Species list for the Coastal Carolinas prioritization.

## ACKNOWLEDGMENTS

We are grateful to the Knoblock Family Foundation, J. Ellis, B. Doolin, and E. Doolin for their support of this work. We are also extremely thankful to the many data contributors who graciously shared their tracking data with Audubon. Please see Table S2 for the complete list of data contributors. The USGS Bird Banding Lab was also kind enough to share band reencounter data with us for this work. We greatly appreciate the time invested by over 150 of Audubon’s fieeld staff and leadership who contributed their knowledge and perspectives to the full annual cycle prioritizations. We thank the many individuals from Audubon North and South Carolina for their contributions to the Coastal Carolinas regional prioritization. We are grateful to Audubon Americas team for discussions leading to the refienement of analyses for the business plan. This material uses data from the eBird Status Project at the Cornell Lab of Ornithology, eBird.org. Any opinions, fiendings, and conclusions or recommendations expressed in this material are those of the author(s) and do not necessarily reflect the views of the Cornell Lab of Ornithology.

## Notes

### Competing Interest Statement

The authors have declared no competing interest.

