## Supplementary material for "A framework for linking hemispheric, full annual cycle prioritizations to local conservation actions for migratory birds": Figure S1. Migratory Connectivity Regions map.

**Figure S1.** Map of administrative units used for the Zonation prioritization. Administrative units were primarily used to ensure that a full range of conservation values were relatively evenly dispersed across the hemisphere. These units were originally used as migratory connectivity regions (MCRs) in Meehan et al. 2022, which was the source of the integrated migration layers used in the full annual cycle prioritizations.

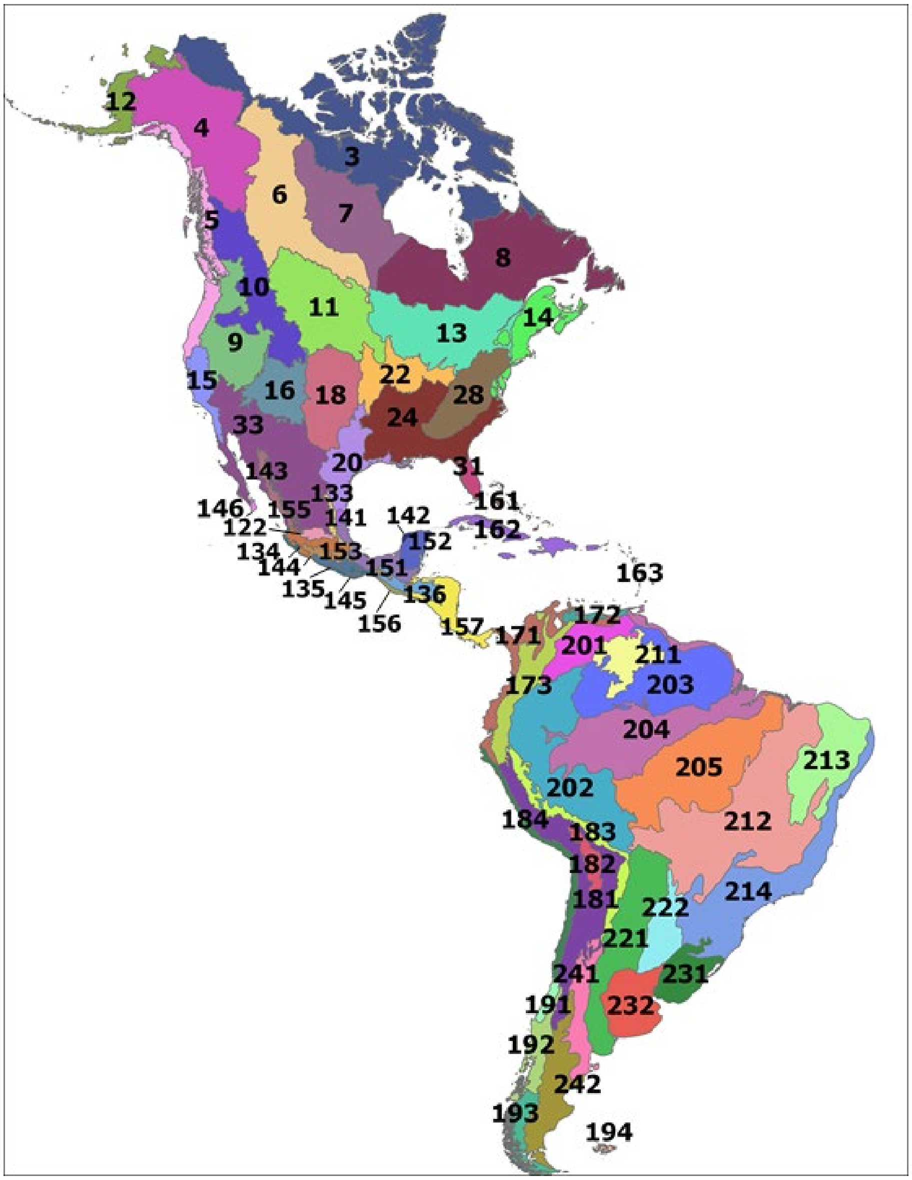
